# A laboratory demand optimisation project in primary care

**DOI:** 10.1101/247163

**Authors:** Magda Bucholc, Maurice O’Kane, Brendan O’Hare, Ciaran Mullan, Paul Cavanagh, Siobhan Ashe, KongFatt Wong-Lin

**Affiliations:** Intelligent Systems Research Centre, University of Ulster, Magee Campus, Londonderry BT48 7JL, Northern Ireland, UK; Altnagelvin Area Hospital, Western Health and Social Care Trust, Glenshane Road, Londonderry BT47 6SB, Northern Ireland, UK; Western Local Commissioning Group, Gransha Park House, Clooney Road, Londonderry BT 47 6FN, Northern Ireland, UK

**Keywords:** laboratory test, test request variability, clinical intervention, clinical pathology, primary care

## Abstract

**Background:** There is evidence of increasing use of laboratory tests with substantial variation between clinical teams which is difficult to justify on clinical grounds. The aim of this project was to assess the effect of a demand optimisation intervention on laboratory test requesting in primary care.

**Methods:** The intervention comprised educational initiatives, feedback to 55 individual practices on test request rates with ranking relative to other practices, and a small financial incentive for practices to engage and reflect on their test requesting activity. Data on test request numbers were collected from the laboratory databases for consecutive 12 month periods; pre‐intervention 2011-12, intervention 2012-13, 2013-14, 2014-15, and post-intervention 2015-16.

**Results:** The intervention was associated with a 3.6% reduction in the mean number of profile test requests between baseline and 2015-16, although this was seen only in rural practices. In both rural and urban practices, there was a significant reduction in-between practice variability in request rates. The mean number of HbA_1c_ requests increased from 1.9 to 3.0 per practice patient with diabetes. Variability in HbA_1c_ request rates increased from 23.8% to 36.6%. At all considered time points, test request rates and variability were higher in rural than in urban areas.

**Conclusions:** The intervention was associated with a reduction in both the volume and between practice variability of profile test requests, with differences noted between rural and urban practices. The increase in HbA_1c_ requests may reflect a more appropriate rate of diabetes monitoring and also the adoption of HbA_1c_ as a diagnostic test.

**Strengths & limitations of the study:**

- We assessed the effect of a laboratory demand optimisation intervention both on the value and between GP practice variability in laboratory test requesting.
- The changes in laboratory test requesting were separately evaluated for rural and urban GP practices.
- Other factors (GP practice organisation, characteristics of general practitioners) potentially affecting between practice differences in laboratory test ordering were not taken into account due to data unavailability.
- The demand management initiative was not accompanied by the cost-effectiveness analysis.
- The demand optimisation intervention was conducted in a Northern Ireland (NI) Western Health and Social Care Trust and the findings have not been independently replicated in any other NI trusts.

## Introduction

Despite the important role of laboratory testing in the diagnosis and monitoring of disease, there is concern about the increasing use of laboratory tests and in particular, the substantial variation in test ordering rates between clinical teams [1]. In the UK laboratory test requests increased by approximately 5% per year in the period 2012-15 [2]. While it is difficult to specify for most tests what an ‘appropriate’ test request rate might be for a given patient population, it is probable that variability in test ordering rates reflects both inappropriate over-and under-requesting [3,4]. Several studies have suggested that around 25-40% of test requests may be unnecessary [5,6], and do not contribute to patient management. This may reflect a lack of knowledge about the appropriate use of individual tests, the use of different clinical guidelines and protocols, inability to access previous results or defensive behaviour of physicians due to fear of errors and medical malpractice litigation [7-10]. Unnecessary testing is not only wasteful of resources but impacts on patients directly through the requirement for venepuncture and the follow up of minor (and possibly insignificant) abnormalities detected and which may cause patient anxiety. Inappropriate under requesting may cause harm through failure to diagnose or manage disease optimally.

Various demand optimisation interventions have been proposed to encourage more appropriate laboratory testing and include educational initiatives on the role and limitations of individual tests and appropriate retest intervals [11-14], [15], feedback on test usage [16-19], redesigning of laboratory tests request forms [20], the introduction of locally agreed clinical guidelines [21,22] and prompts on electronic test ordering systems. The effectiveness of such interventions is variable and depends in part on local factors and local clinical team engagement. Furthermore, such interventions may be time consuming and expensive; a study on educational interventions conducted in hospital settings showed that the savings on the direct hospital costs resulting from interventions were smaller than the cost of interventions [23].

The aim of this study was to investigate the effect of a laboratory demand optimisation intervention in a primary care setting on both laboratory test request rates and on the variability between practices in test request rates.

## Materials and methods

This study was undertaken in 55 separate primary care medical practices within the catchment area of the Western Health and Social Care Trust (WHSCT). The WHSCT provides laboratory services to these practices with networked laboratories in each of the three large urban centres of Londonderry, Omagh, and Enniskillen. The patient population served by the 55 practices over the 5-year study period was 316 382 (2011-12), 316 688 (2012-13), 318 057 (2013-14), 319 383 (2014-2015), and 326 429 (2015-2016).

The primary care practices were situated in both rural and urban areas using data from the Census Office of the Northern Ireland Statistics and Research Agency [24]. Since the Northern Ireland settlement classification does not give continuous spans of particular area types, a practice was designated as urban if its postal address was situated in a settlement of more than 10,000 residents following the urban-rural classification thresholds used by the Department for Environment, Food and Rural Affairs (DEFRA) and the Department for Communities and Local Government (DCLG) [24]. Under this definition, 31 practices were designated as urban and 24 as rural.

Data on laboratory test requests from individual primary medical practices were studied over five consecutive 12 month periods (1 April to 31 March) from 2011-12 (the pre‐intervention or ‘baseline’ period) to 2015-16. Test request data were extracted from the laboratory databases of the Altnagelvin Area Hospital, Tyrone County Hospital, and the Erne Hospital (subsequently the South West Acute Hospital). Information on individual primary care practices regarding registered patient numbers, the number of male patients, and patients with diabetes was obtained from the Western Health and Social Services Board Integrated Care Partnership.

The following test groups were studied: electrolyte profile, lipid profile, thyroid profile (FT4 and TSH), liver profile, immunoglobulin profile, glycosylated haemoglobin (HbA_1c_), and prostate-specific antigen (PSA). The number of profile tests (electrolyte profile, lipid profile, thyroid profile, liver profile, immunoglobulin profile) requested in each practice was standardised against the number of registered patients in the practice and expressed as requests per 1000 patients. HbA_1c_ was standardised against the number of patients with diabetes per practice while PSA was standardised against the number of male patients per practice.

Throughout the study period laboratory requests from primary care were ordered on a paper laboratory request form. All of the test considered here (with the exception of immunoglobulins) were listed on the request form and could be ordered by ticking a box on the test request form adjacent to the test profile name; an immunoglobulin profile was ordered by free text entry on the request form.

Test requesting rates were studied before and after a three year intervention designed to support optimal use of laboratory testing. The intervention package was developed in conjunction with the Western Local Commissioning Group (responsible for commissioning and managing primary care services and which included senior primary care doctors). The intervention included several discrete elements. Firstly, awareness of the intervention was promoted through educational sessions on the benefits to patients and clinical teams of optimal use of laboratory tests. Secondly, educational material was developed in conjunction with primary care clinicians which covered the major clinical indications for a range of considered requested tests and appropriate retest intervals. This was distributed to all primary care teams and was supplemented by presentations at educational meetings. Thirdly, all primary care teams were asked to engage in the process of reviewing test requesting procedures within their practice, and to reflect on the information provided on their practice test requesting rates and ranking in comparison to other practices. The active intervention took place over the three year period: 2012-13, 2013-14, and 2014-15.

Prior to the intervention each practice received information on its standardised test request rates (see below) over the previous year (baseline period) and its ranking in relation to standardised test request rates of all other practices served by the laboratory.

The Western Local Commissioning Group (WLCG) made available funding to incentivise participation in this process. All participating primary care practices received a payment of £0.30 per patient registered on their practice list to engage in the process or reviewing and reflecting on test requesting activity. Changes in the absolute numbers of standardised test requests and between-practice variability in standardised test request rates were compared to the pre‐intervention (‘baseline’) period (April 2011 – March 2012).

Variability between practices in standardised test request rates was expressed as coefficient of variation (CV) whereas differences in the variance between pre‐ and post-intervention period were tested using the Bonett-Seier test [25]. A paired t-test was used to compare mean numbers of laboratory test requests from pre- and post-intervention period.

Spearman’s rank correlation was used to study relationships between standardised requesting rates for three of the most commonly request tests (electrolyte, liver, and lipid tests) within individual practices [26]. All statistical analyses were performed using R statistical software, version 3.3.3 (R Foundation for Statistical Computing, Vienna, Austria).

## Results

The total number of profile test requests for all practices fell from a mean of 1554 per 1000 patients pre‐intervention to 1498 per 1000 patients one year post intervention (a reduction of 3.7%; *p* = 0.09) (Table 1). Rural practices had a higher average standardised profile request rate than urban practices at all time points: baseline, during the intervention and at one year post intervention. However, the reduction in the mean number of test requests was seen exclusively in rural practices where requests fell by 9% (*p* = 0.01) as compared with no significant change in urban practices.

**Table 1.**
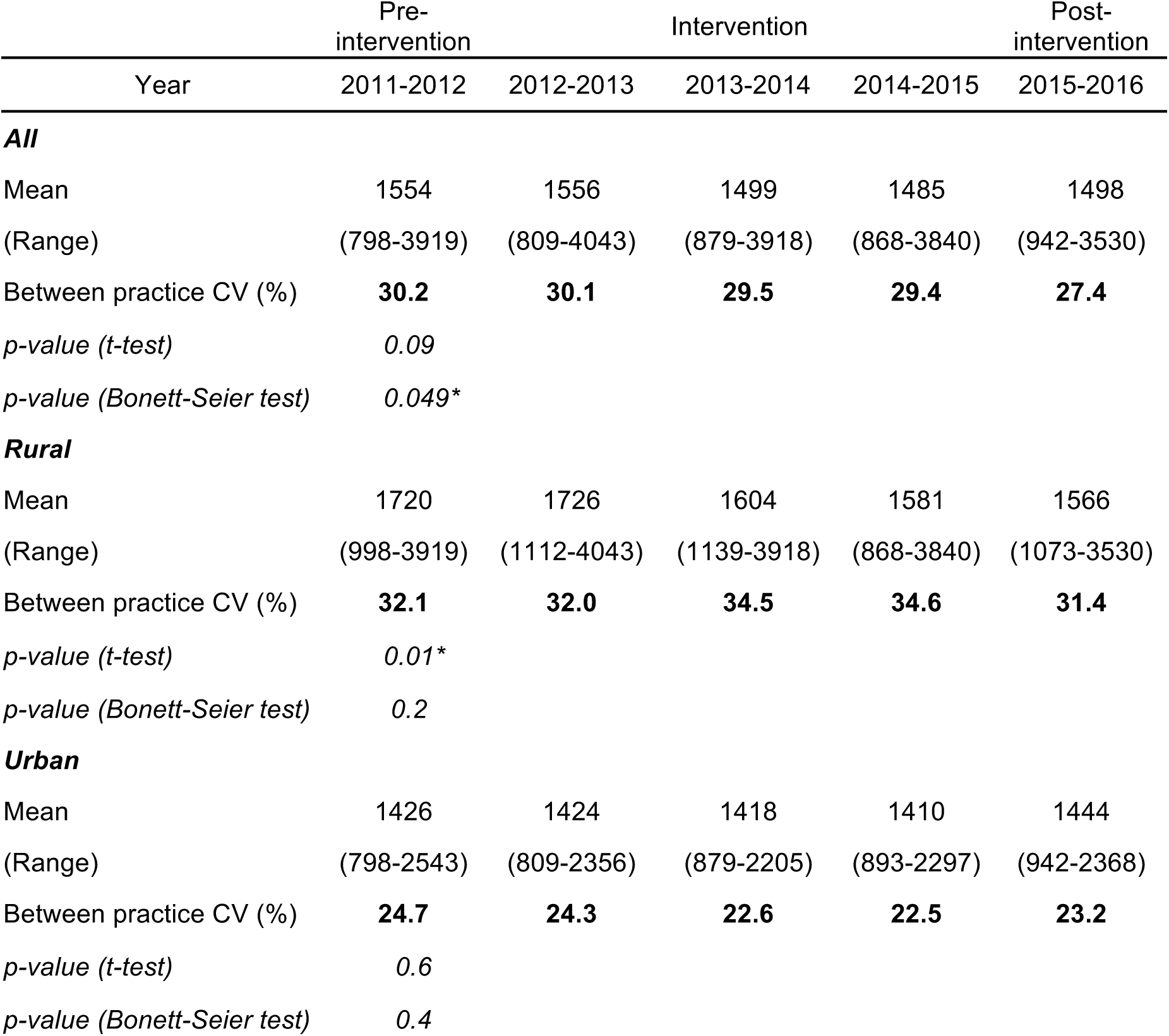
Standardised profile test request rates per 1 000 patients pre‐ and post-intervention for all practices combined and for rural and urban practices. T-test p-value refers to the significance level evaluating differences between the mean number of request rates in pre‐ and post-intervention period. The p-value of Bonett-Seier test refers to the significance level assessing the difference in variances in pre‐and post-intervention period. Asterisk: Statistically significant difference (*p* < 0.05) between the pre‐ and post-intervention data.

|  | Pre-intervention |  | Intervention |  | Post-intervention |
| --- | --- | --- | --- | --- | --- |
| Year | 2011-2012 | 2012-2013 | 2013-2014 | 2014-2015 | 2015-2016 |
| **All** |  |  |  |  |  |
| Mean | 1554 | 1556 | 1499 | 1485 | 1498 |
| (Range) | (798-3919) | (809-4043) | (879-3918) | (868-3840) | (942-3530) |
| Between practice CV (%) | **30.2** | **30.1** | **29.5** | **29.4** | **27.4** |
| *p-value (t-test)* | 0.09 |  |  |  |  |
| *p-value (Bonett-Seier test)* | 0.049* |  |  |  |  |
| **Rural** |  |  |  |  |  |
| Mean | 1720 | 1726 | 1604 | 1581 | 1566 |
| (Range) | (998-3919) | (1112-4043) | (1139-3918) | (868-3840) | (1073-3530) |
| Between practice CV (%) | **32.1** | **32.0** | **34.5** | **34.6** | **31.4** |
| *p-value (t-test)* | 0.01* |  |  |  |  |
| *p-value (Bonett-Seier test)* | 0.2 |  |  |  |  |
| **Urban** |  |  |  |  |  |
| Mean | 1426 | 1424 | 1418 | 1410 | 1444 |
| (Range) | (798-2543) | (809-2356) | (879-2205) | (893-2297) | (942-2368) |
| Between practice CV (%) | **24.7** | **24.3** | **22.6** | **22.5** | **23.2** |
| *p-value (t-test)* | 0.6 |  |  |  |  |
| *p-value (Bonett-Seier test)* | 0.4 |  |  |  |  |

The between practice coefficient of variation for profile test requests fell from 30.2 % pre‐intervention to 27.4% one year post intervention (*p* = 0.049). Rural practices had a higher between practice coefficient of variation than urban practices at all time points (Table 1). There was no significant difference between urban and rural practices in the number of registered patients per general practitioner.

For HbA_1c_, there was an increase in mean test request rates from 1.9 requests per patient with diabetes pre‐intervention to 3.0 diabetes post-intervention (*p* =0.00001) (Table 2). Variability for HbA_1c_ increased from 23.8% to 36.6% (*p* = 0.00001). The statistically significant increase in variation for HbA_1c_ was observed both in rural (*p* = 0.00031) and urban (*p* = 0.008) areas (Table 2).

**Table 2.**
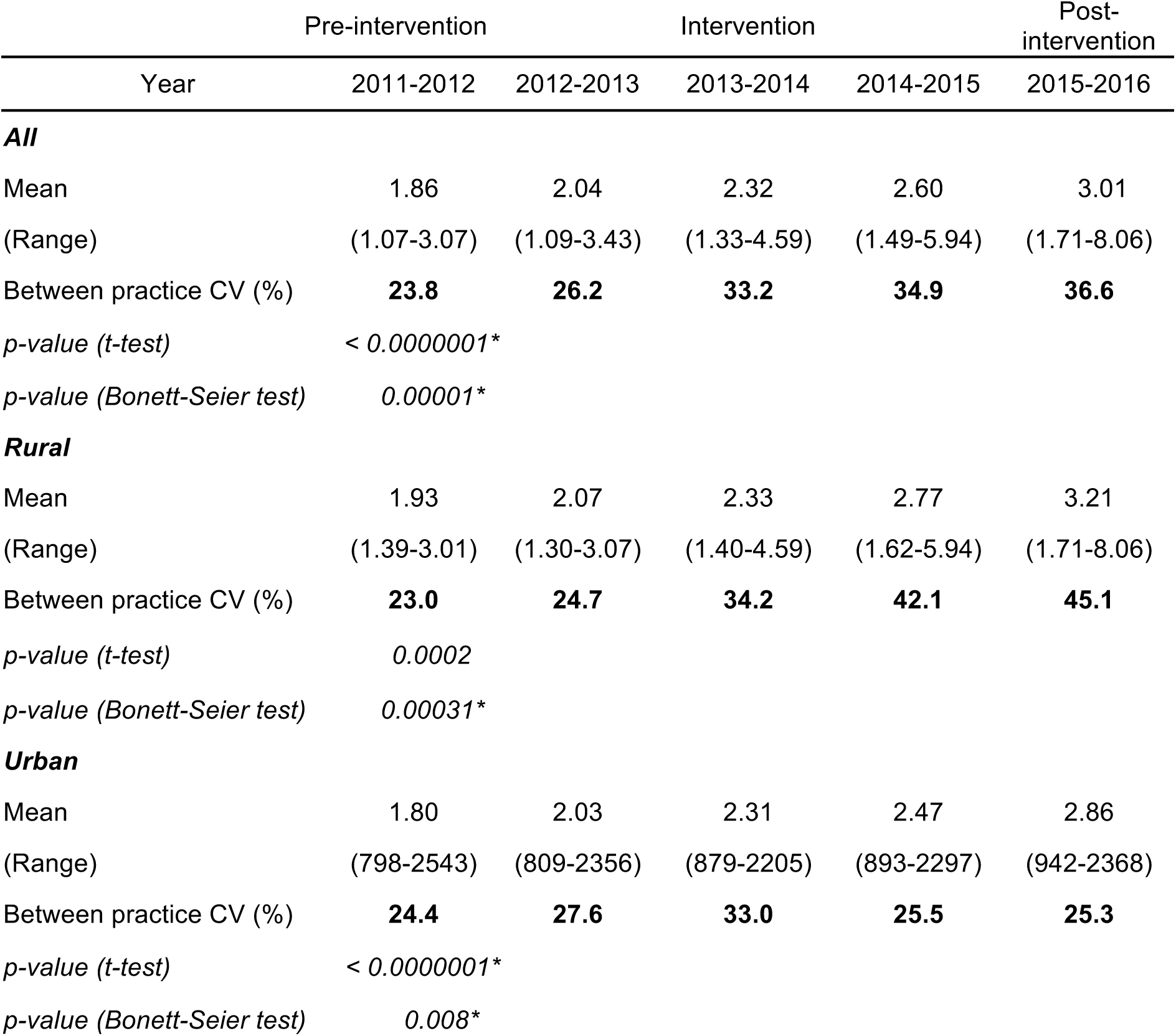
Standardised test request rates pre‐ and post-intervention for HbA_1c_(expressed as number of tests per patient with diabetes). Asterisk: Statistically significant difference (*p* < 0.05) between the pre‐ and post-intervention data.

The mean number of PSA requests per 1000 male patients increased from 69.4 to 82.9 following the intervention (*p* = 0.006) (Table 3). However, there was no significant change in between practice variability.

Finally, there were high correlations within practices for individual profile test types: electrolyte and liver profiles (R = 0.83), and lipids and liver profiles (R=0.67) (Figure 1).

**Figure 1.**
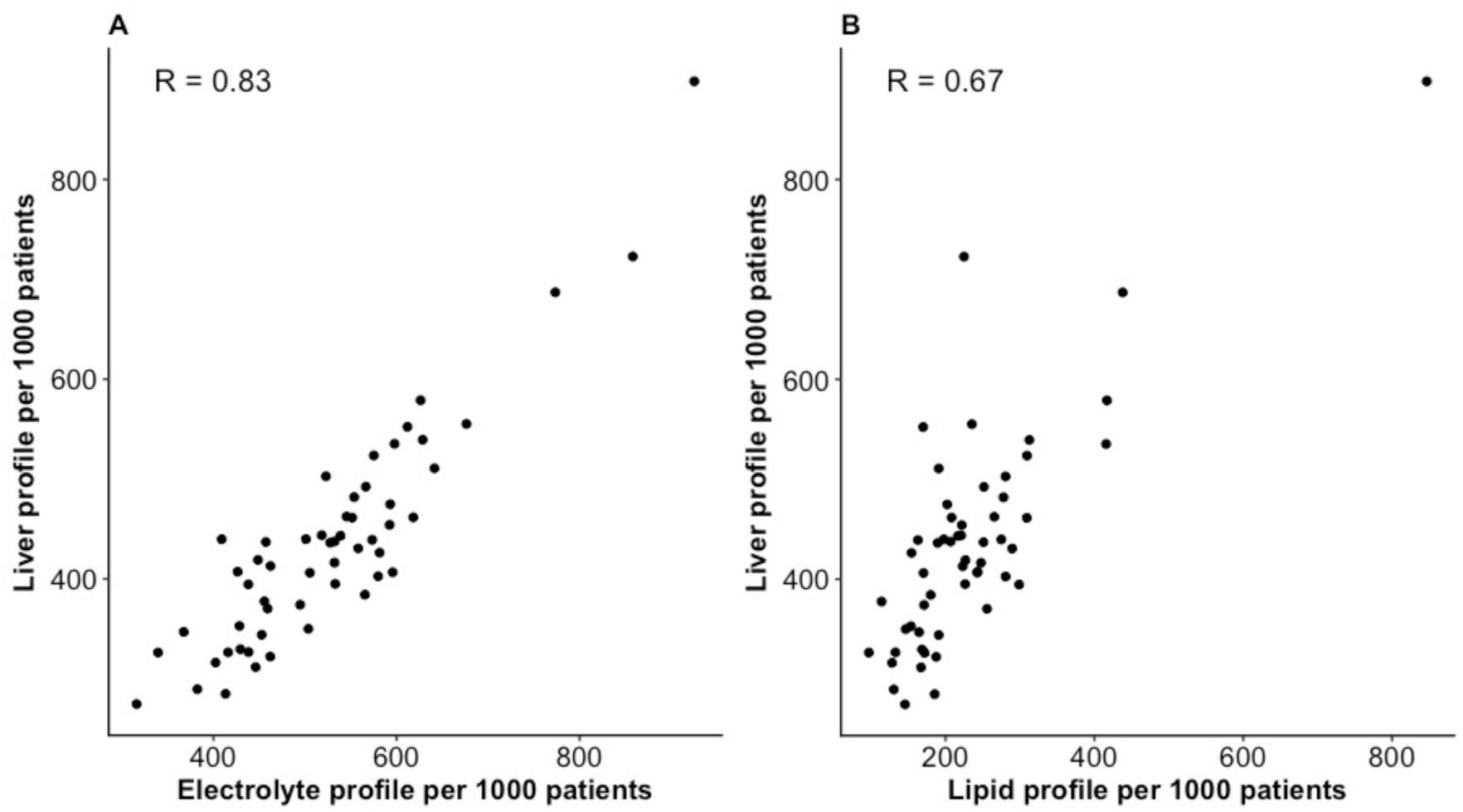
Practice standardized requests for electrolyte profiles plotted against liver profiles (A), and lipid profiles plotted against liver profiles (B), with Spearman’s rank correlation coefficients (R).

**Table 3.**
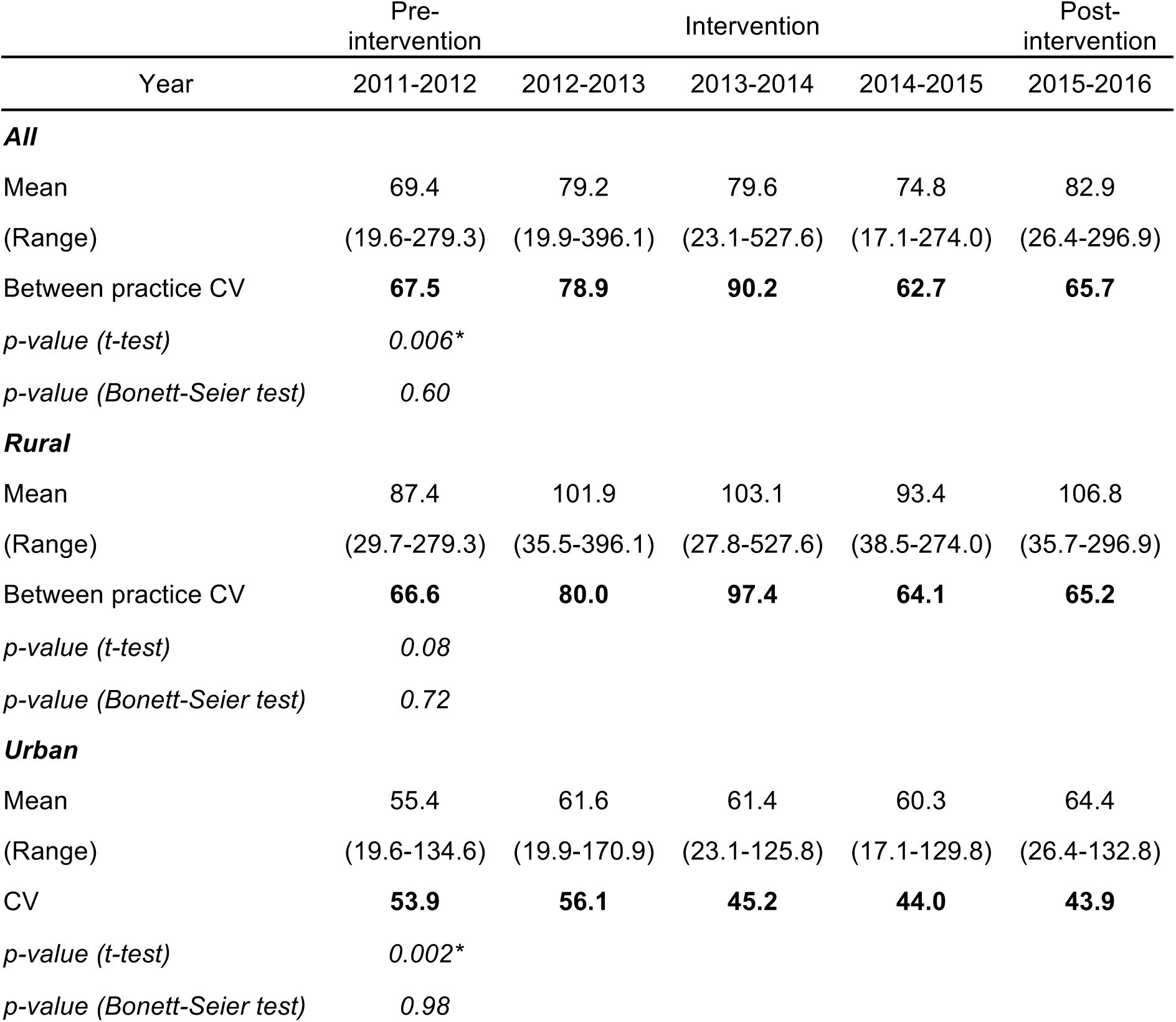
Standardised test request rates pre‐ and post-intervention for PSA(expressed as number of tests per 1 000 male patients). Asterisk: Statistically significant difference (*p* < 0.05) between the pre‐ and post-intervention data.

## Discussion

While it may be challenging to define what represents an appropriate rate of requesting for most tests, it is certainly difficult to justify very high levels of variability between clinical teams providing care to broadly similar groups of patients within a single healthcare system. This study found high levels of baseline variability between primary care practices in standardised biochemistry profile test request rates both in rural (32.1%) and urban areas (24.7%). There is little reason to believe that there were significant differences in the characteristics of the practice patient populations within each of rural and urban areas in terms of disease prevalence or morbidity that might explain such high variability. It is therefore likely that the variability observed reflects differing behaviours and perceptions between clinical teams as to the value and role of individual tests in patient assessment.

The baseline standardised profile test request rates were significantly higher in rural than in urban practices. The reasons for this are unclear and were beyond the scope of investigation of the present study. However possible explanations include differences in practice organisation and workflow, differences in the characteristics of general practitioners such as training, background, age which might lead to differences in approach to patient assessment and testing [27,28].

The intervention employed to optimise demand was associated with two effects. Firstly, there was a reduction of 3.7% in total standardised profile test requests (as measured at one year post intervention). However, this was accounted for entirely by a reduction in rural practices. Secondly, there was also a significant reduction in between practice variability in test requesting from 30.2% to 27.4% and this was seen in both urban and rural practices. During and post-intervention, the standardised test request rates and variability continued to be higher in rural than urban practices.

For HbA_1c_ the standardised test rate per patient with diabetes increased from 1.9 to 3 tests per patient with diabetes per year. Best practice guidelines suggest measuring HbA_1c_ two to three times per year in patients with diabetes and this had been highlighted in the educational material that formed part of the intervention [29]. The increased testing rate may therefore reflect more appropriate monitoring of patients with diabetes. However, as it was not possible to distinguish HbA_1c_ samples which had been requested for diabetes monitoring from those requested for the purposes of diabetes diagnosis, it is difficult to be certain. The use of HbA_1c_ as a diagnostic test for diabetes mellitus had been introduced in 2012 i.e. during the baseline period and it is possible that the increase in requesting (and the observed increase in between practice variability) reflected its adoption as a diagnostic test rather than as a monitoring test.

Within individual practices, there was high correlation between standardised request rates for different test profiles. The reasons for such correlations are unclear. In some instances, there may be good reasons why different test profiles should be requested together e.g. monitoring liver enzymes along with lipids for patients on statin therapy. In other cases, there may be patients with complex medical conditions and a number of co-morbidities in whom it is appropriate to request a number of test profiles simultaneously. A further possibility is that the co-requesting of different test profiles reflects a ‘scatter gun’ approach to test requesting. This may also have inadvertently been facilitated by the design of the test request form on which tests are requested simply by ticking a box beside the relevant test profile.

Previous studies on demand optimisation in primary care have yielded varying results with some studies showing reductions of up to 12% [17,19,30-32] following a range of educational and feedback interventions or guideline driven decision support systems. Studies which targeted the utilization of specific laboratory tests also showed that the interventions generally produced changes in the desired direction. For example, educational initiatives were found to improve significantly the management of albuminuria [33], oral anticoagulation [34], C-reactive protein [35], HbA_1c_ [36-38], lipids [36-39], and Pap testing [40]. The improvement was more likely to be observed when more than one type of intervention was used at a time [38,41].

Although numerous previous studies had documented high degrees of variability in test requesting between primary care teams [32,42,43], a unique feature of the present study was that it assessed the effect of the intervention on between practice variability in test requesting. The reduction in variability found here suggests that the intervention was associated with a more standardised approach to patient investigation and monitoring.

## Conclusions

In conclusion, the demand optimisation intervention undertaken here showed a small but significant reduction in reducing unwarranted variability between practices in test requesting rates.

## Data

The information on datasets supporting this article have been provided in the supplementary material.

## Competing interests

We have no competing interests

## Authors’ contributions

Contributors: MB performed the analysis and interpretation of the results, and wrote the manuscript. She is guarantor. MJO, BOH, CM and PC designed and carried out the demand management intervention. MJO wrote the manuscript. BOH, CM and PC edited the manuscript. SA monitored the data collection and edited the manuscript. KWL initiated the collaborative project, guided the data analysis and interpretation of the results, and wrote the manuscript.

## Funding

This work was performed under the Northern Ireland International Health Analytics Centre (IHAC) collaborative network project funded by Invest NI through Northern Ireland Science Park (Catalyst Inc). The funder had no role in study design, data collection and analysis, decision to publish, or preparation of the manuscript.

## Acknowledgments

The authors would wish to thank the IHAC collaborative network, especially Colm Hayden, Brendan Bunting and Le Roy Dowey for helpful discussions; Graham Moore, Austin Tanney, and Paul Barber for computing and technical support; and Stephen Lusty and Peter Devine for administrative support.

